# Rapid acquisition of near optimal stopping using Bayesian by-products

**DOI:** 10.1101/844969

**Authors:** Joshua Calder-Travis, Emma Slade

**Affiliations:** Department of Experimental Psychology, University of Oxford, Radcliffe Observatory Quarter, Woodstock Road, Oxford, OX2 6GG, United Kingdom; Rudolf Peierls Centre for Theoretical Physics, University of Oxford, Clarendon Laboratory, Parks Road, Oxford OX1 3PU, United Kingdom

## Abstract

Prior to any decision, animals (including humans) must stop deliberating. Both accumulating evidence for too little or too long can be costly. In contrast to accounts of decision making, accounts of stopping do not typically claim that animals use Bayesian posteriors. Considering a generic perceptual decision making task we show that, under approximation, only two variables are relevant to the question of when to stop evidence accumulation; time and the maximum posterior probability. We explored the rate at which stopping rules are learned using deep neural networks as model learners. A network which only used time and the maximum posterior probability learned faster than any other network considered. Therefore, such an approach may be highly adaptive, and animals may be able to reuse the same neural machinery they use for decisions for stopping. These results suggest that Bayesian inference may be even more important for animals than previously thought.

## 1 Introduction

Prior to any decision, animals (including humans) must terminate deliberation. In perceptual decisions, the longer an animal waits, the more sensory information they can gather, and hence the more likely their decision is to be correct (Ratcliff & Rouder, 1998). However, waiting is itself costly. For example, the longer a seal waits, the more likely it is to correctly identify the fish in the distance. On the other hand, the longer it waits, the more likely it is that the fish (whether predator or prey) will spot them. Hence, stopping rules, not just decision rules, are important to survival.

The idea that animals use optimal decision rules has been studied extensively. Bayes rule provides the mathematical formalism that can be used to make inferences about the environment from neural signals generated by sense organs (Ma & Jazayeri, 2014). With a posterior distribution over the state of the environment, animals can take actions which maximise the expected reward. The idea that we use Bayesian inference to make decisions has received support in domains as diverse as perceptual decisions (Tassinari, Hudson & Landy, 2006), multi-sensory decisions (Ernst & Banks, 2002), and word recognition (Norris, 2006). Additionally, we have several accounts of how the brain might represent and compute with probability distributions (Ma, Beck, Latham & Pouget, 2006; Sahani, 2003; Bogacz & Gurney, 2007).

Many models of stopping instead involve rules which are not optimal, and do not use posterior probabilities over states of the environment. These models typically assume that each of the *N* options in a decision making task generate a noisy signal in the brain, representing the “evidence” for that option (Bogacz, Brown, Moehlis, Holmes & Cohen, 2006; Usher & McClelland, 2001). The signal which corresponds to the correct alternative has a higher mean. These evidence signals are then accumulated in some form, as the accumulation of these noisy signals reduces the effects of noise. Models often assume that animals use a non-optimal stopping rule, such as the difference between accumulators (e.g. diffusion decision model; Ratcliff and McKoon, 2008; optimal in some cases, Gold and Shadlen, 2001, 2002; Moran, 2015; Wald and Wolfowitz, 1948), the maximum accumulator (e.g. race model; Vickers, 1970; Bogacz et al., 2006), leaky accumulators and/or accumulators which inhibit each other to some extent (Usher & McClelland, 2001).

The multihypothesis sequential probability ratio test (MSPRT) is an interesting exception. The MSPRT has been used to model the stopping rules employed by animals when deciding between multiple (*N* > 2) options, and unlike other models applicable to this situation, it claims that animals use posterior probabilities for stopping (Bogacz & Gurney, 2007; Baum & Veeravalli, 1994). While this model uses posterior probabilities for stopping, it is only optimal under conditions in which animals make very few errors, or the cost of errors are very small (Dragalin, Tartakovsky & Veeravalli, 1999).

The optimal stopping rule for animals facing certain kinds of perceptual and value-based decisions has been derived (Drugowitsch, Moreno-Bote, Churchland, Shadlen & Pouget, 2012; Tajima, Drugowitsch & Pouget, 2016; Tajima, Drugowitsch, Patel & Pouget, 2019). In the *N* -dimensional space defined by the *N* evidence accumulators, the optimal stopping policy is to terminate deliberation when the point in this space which represents the state of the *N* accumulators, reaches a time-dependent curved bound (Tajima et al., 2019; Fig. 1). Calculating this bound involves first calculating the value function, which gives the reward expected in the future, if one behaves optimally. Unfortunately, computing the value function involves considering some distant future time, and then working backwards, calculating the value function time step by time step (Drugowitsch et al., 2012).

**Figure 1:**
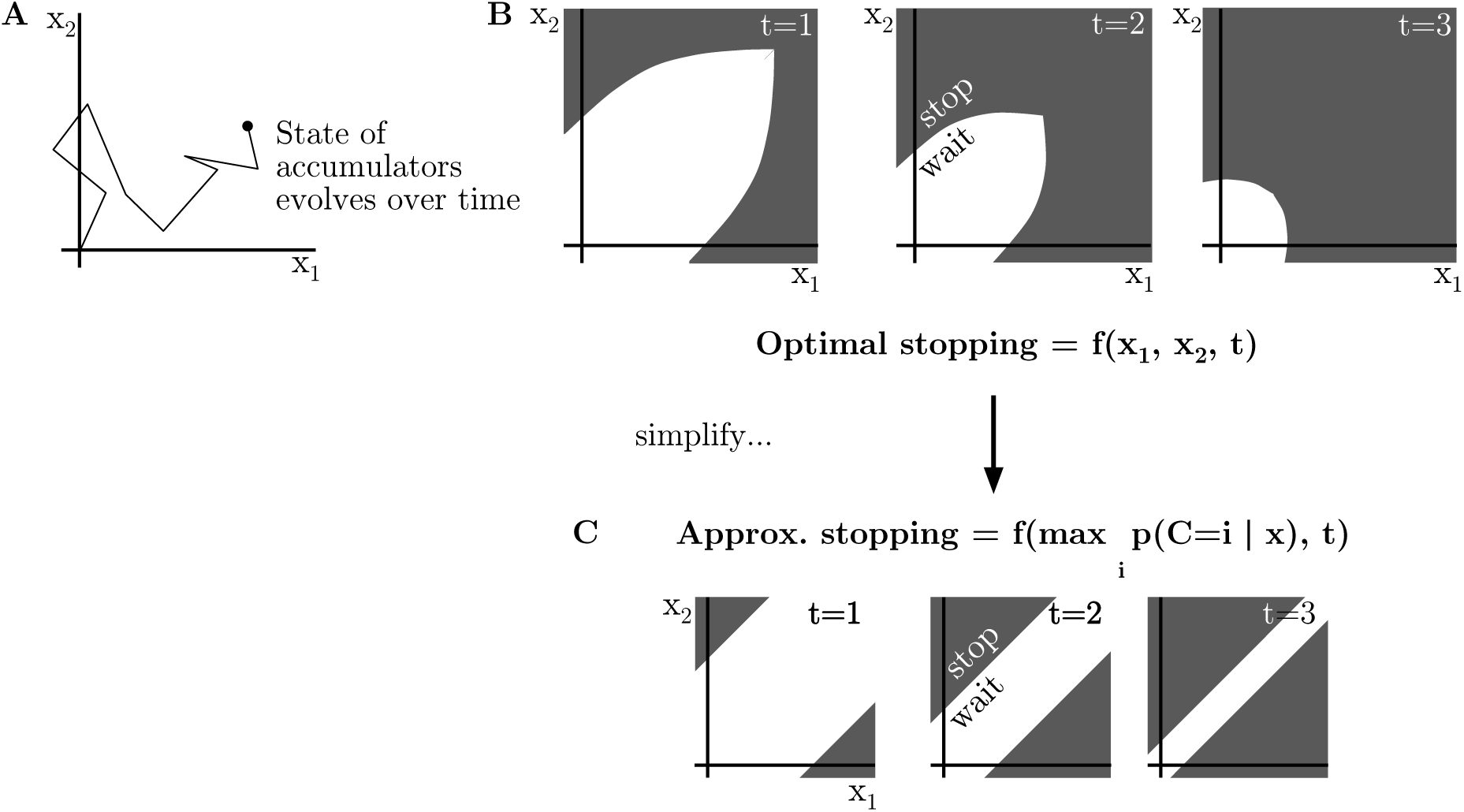
Animals may use *N* accumulators to track the cumulative evidence for the *N* choices. The case of *N* = 2 is shown here. (A) The state of the accumulators can be represented as a point in *N* dimensional space that moves over time. Each dimension represents the value of one accumulator *x*_*i*_. (B) The optimal stopping rule involves accumulating until a criterion on time, *t*, and the state of the *N* accumulators, *x*_*i*_, is met (Tajima, Drugowitsch, Patel & Pouget, 2019). Criterion shown only for purposes of illustration, and not derived. (C) It may be possible to find a stopping rule which approximates the optimal stopping rule, but which uses a criterion on fewer variables.

The computational intensity of computing the optimal stopping rule makes it less plausible that animals compute the exact rule. An alternative is that animals gradually learn a mapping from the value of the accumulators and elapsed time, to stopping vs. waiting. Unfortunately, this also likely to be challenging, especially when the number of choice options is high. In this case the animal must learn to map from a high dimensional vector (with one entry for each accumulator state). Moreover, the optimal stopping rule will change if task or environmental statistics change (Drugowitsch et al., 2012). Hence, in naturalistic settings, learning optimal stopping rules will only be feasible if learning is rapid.

In groundbreaking work, Tajima et al. (2019) provided a potential solution to these issues by showing that approximately optimal stopping rules may be considerably simpler than optimal stopping rules. Specifically, they showed that if the state of the accumulators is mapped onto a curved manifold, approximately optimal stopping can be achieved by stopping when the value of the maximum mapped accumulator crosses a threshold. This work leaves open a number of questions. Could the brain recycle representations of posterior probabilities (which are non-linear transformations of the accumulators), saving the need for additional neural hardware? How robust is the mapping used by Tajima et al. (2019); must this transformation be adapted to task and environmental conditions?

Here we consider a generic perceptual decision making task and show that, under a wide range of conditions, a criterion on time and the maximum posterior probability, rather than a criterion using time and all *N* accumulator values, is sufficient for approximately optimal stopping (Fig. 1). Hence, when learning an optimal stopping rule, animals only need to learn a mapping from two variables, to waiting vs. stopping. Additionally, we find that this approximation facilitates rapid learning of the stopping rule. Taken together, these results suggest that the brain can recycle Bayesian posterior probabilities computed for the purposes of optimal deciding, for the purposes of approximately optimal stopping.

In Sec. 2 we discuss in detail the mathematical formalism. Specifically, we outline how one may approximate optimal stopping using a criterion on time and maximum posterior probability. Apart from the first paragraph, and equations 1 and 16, Sec. 2 may be safely skipped by a reader uninterested in the derivation of this result. In Sec. 3 we discuss the computational model learners we use to test our hypothesis, and discuss the results using the model learners in Sec. 4. Finally in Sec. 5 we conclude and outline possible areas for future work.

## 2 Mathematical results

### 2.1 Decision problem and posterior

We formalise a perceptual decision making task using the standard idea that, for each option, the brain computes a noisy evidence signal (Bogacz et al., 2006). The goal of an observer is to determine which object or property, *C*, is the cause of the observed evidence samples *δ****x***(1), *δ****x***(2), …***x***(*M*). *δ****x***(*t*) denotes a vector containing the values of the evidence signals for each of the *N* options, at time step *t*. The prior probability of object *C* = *i* is *p*(*C* = *i*), and *C* = *i* generates average evidence samples given by ***µ***(*i*). ***µ***(*i*) indicates a vector, the same length as the number of possible values of *C*, where each element is given by

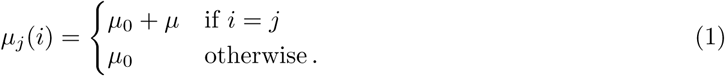

Every time step the object generates associated evidence samples, *δ****x***(*t*), according to

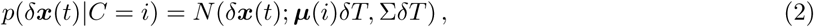

where *t* indicates the current time step, *δT* is the duration of a time step, *N* () indicates the multivariate normal distribution, and Σ is the covariance of evidence samples (Tajima et al., 2019). In supplementary file S1 we find the posterior over *C* using Bayes rule to be

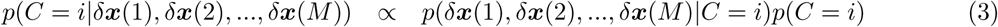

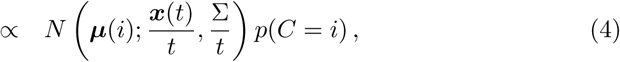

where ***x***(*t*) = ∑_*i*_ *δ****x***(*i*), *t* = ∑_*i*_ *δT* (Drugowitsch et al., 2012).

We consider only covariance matrices which can be written in the following form

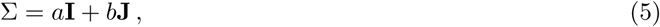

where **J** is a matrix of ones. In supplementary file S1 we find that in this case the posterior becomes

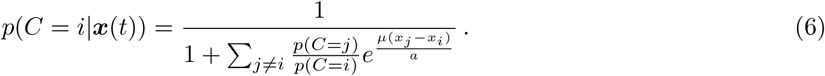

### 2.2 Optimal stopping

If a Bayesian observer stops, they will make the decision which maximises expected reward (Ma & Jazayeri, 2014). If the reward for all options is the same, they will pick the most likely option, i.e. they will have the response, *Ĉ*,

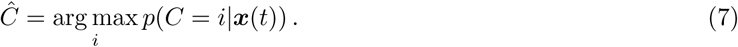

If the observer receives reward *r*_*c*_ for a correct response and *r*_*e*_ for an incorrect response, then the expected reward from deciding, *y*, is

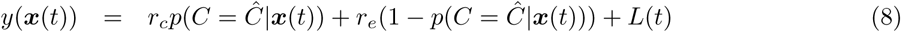

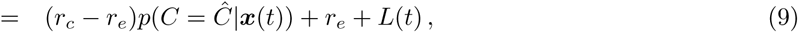

where *L*(*t*) is a function describing the cumulative cost of time. Therefore we can compute the expected reward from deciding. It is more difficult to calculate the expected reward from waiting. Imagine we have a stopping rule, *S*, which describes for which values of ***x***(*t*), and time steps *t* we will stop and decide. Then the expected reward from following *S* from time step *t* onward is

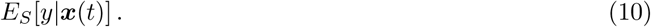

In words, this expression denotes the expected reward, following stopping rule *S*, given that we are currently at ***x***(*t*). An optimal observer will only stop and decide now if there is no way of increasing the expected reward. That is, they will stop now if there is no stopping rule which, if followed from this point on, would lead to an expected reward which is greater than the reward for stopping now (Ferguson, 2011; Øksendal, 2013). The optimal observer will therefore only stop if

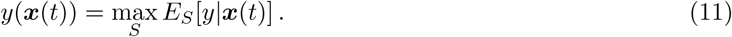

It is difficult to find the set of points which satisfy this equation because is it very difficult to calculate the maximum possible reward under any stopping rule.

### 2.3 Approximating the optimal solution

One reason why it is so difficult to calculate the point at which the expected reward from deciding *y*(***x***(*t*)) equals the maximum expected reward under any stopping rule, is because the future evolution of ***x***(*t*) is unknown, but described by

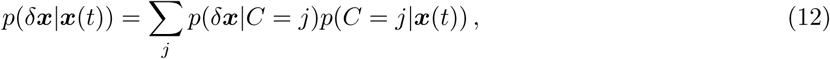

where *δ****x*** is the change in ***x*** at any future time step. Hence, we have to consider the expectation over all possible trajectories, and the time and place at which these paths are terminated according to the stopping rule, to determine the expected reward of waiting.

One approach to finding an approximate solution would be to use some form of myopic approximation and only look ahead a few steps. We make a different approximation here, looking as far ahead as needed, but at each time step only considering the most likely value for *δ****x***. Considering only one value of ***x***(*t*) at each time step reduces the problem of picking a stopping policy to the problem of picking a time to stop.

In supplementary file S1 we consider any two points ***x***^(*d*)^(*t*) and ***x***^(*e*)^(*t*) for which the posterior probability of the most probable option is the same. That is,

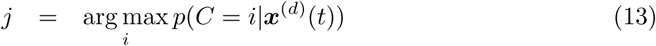

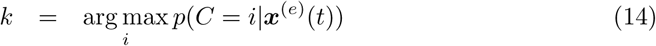

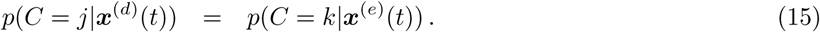

Under the approximation, the expected reward at any later time step *t* + *l* is the same. As the expected future rewards are the same for all stopping times, and only stopping time matters (because ***x*** takes a single path), if it is optimal to stop (or continue) at *t*, ***x***^(*d*)^, then it is also optimal to stop (or continue) at *t*, ***x***^(*e*)^. As ***x***^(*d*)^ and ***x***^(*e*)^ can be any two points for which the posterior probability of the most probable option is the same, we can say that at a time step, *t*, for all points, ***x***, with same maximum posterior probability, it is either optimal to stop at all of them, or none of them. Hence, according to the approximate optimal stopping rule, only two variables are needed to determine whether or not to stop; time and the maximum posterior probability.

## 3 Modelling methods

Having found that an approximately optimal stopping rule exists which uses a stopping criterion on only time and the maximum posterior probability, we set out to test whether learning stopping rules with this simpler form is faster. To do this we use neural networks as model learners.

### Task

The networks were trained to perform a simulated version of the perceptual decision making task described in the previous section (see equations 1 and 16). We used the settings shown in Table 1. We considered every combination of number of choices (2, 4, 10), uncorrelated and correlated accumulators (*b* = 0 and *b* = 0.9), and cost of time (constant, increasing). If constant, the cost of time was 1 per time step, if increasing, the cost of time was 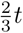, where *t* is the time step. Each trial was made up of 40 time steps, and the networks received 3000 simulated trials (which were then split into training and validation sets).

**Table 1:** Properties of the simulated perceptual decision making task.

| Variable | Expression | Simulation value |
| --- | --- | --- |
| Number of choices | $k$ | 2, 4, 10 |
| Correct mean | $\mu_0 + \mu$ | 1.3 |
| Incorrect mean | $\mu_0$ | 1 |
| Covariance | $b$ | 0, 0.9 |
| Variance | $a + b$ | 1 |
| Correct reward | $r_c$ | 50 |
| Error reward | $r_e$ | -50 |
| Prior | $p(C = i)$ | $1/k$ |
| Cumulative cost of time | $L(t)$ | see text |

For the case of 2 choices, constant cost of time, and no correlation between accumulators, we computed the expected reward for an optimal observer, allowing comparison of actual performance to optimal performance (derivation provided in supplementary file S2).

### Networks

We used deep neural networks implemented in Keras using the TensorFlow backend. The networks comprised of an input layer, an output layer, and 2 hidden layers (see Fig. 2). Each hidden layer had a number of units equal to the number of choices in the decision task, plus 1 (*k* + 1). The output layer contained a single unit.

**Figure 2:**
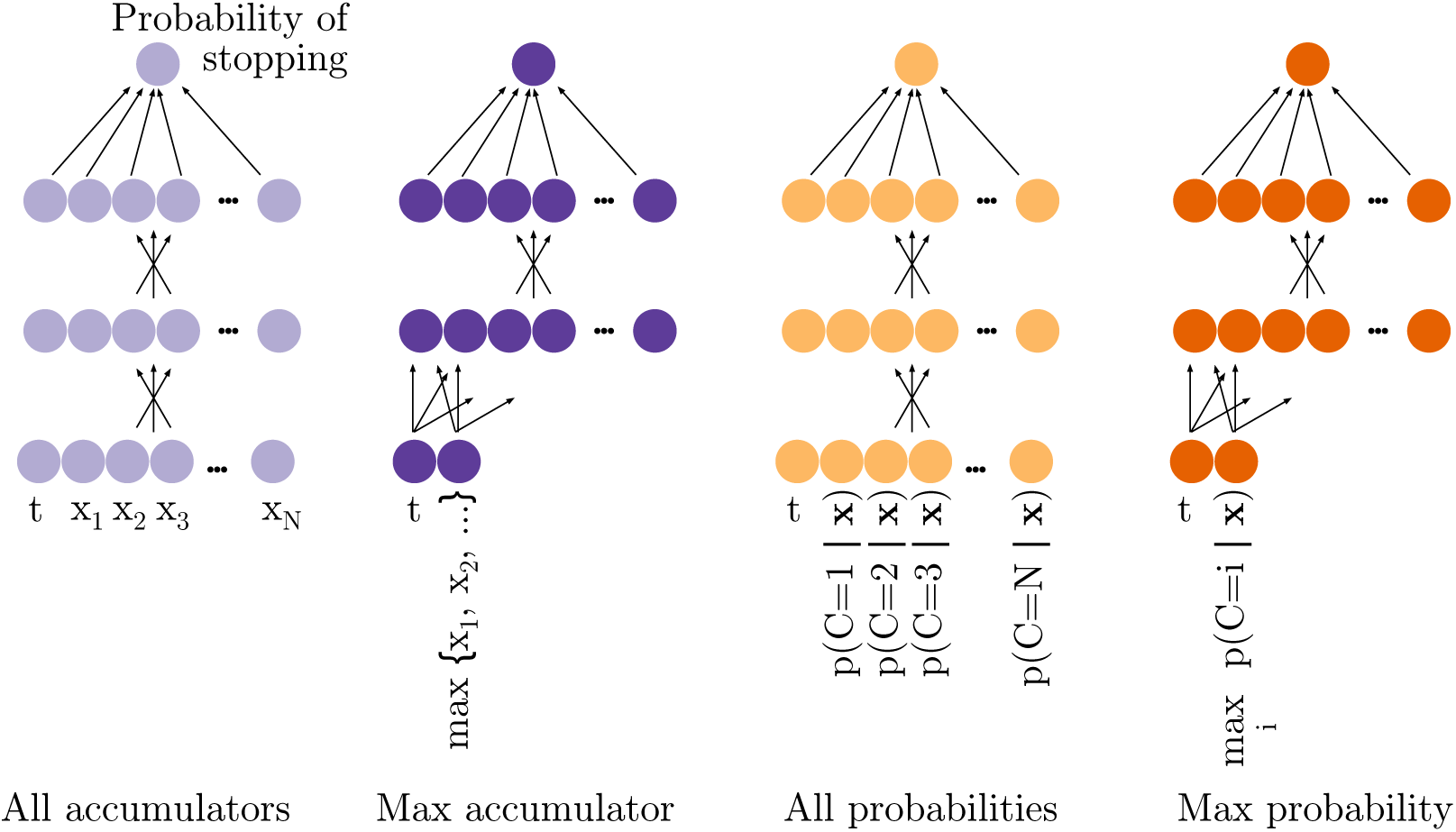
The networks used.

The nature of the input data depended on the network. All networks received the current time step as an input, but varied in their additional inputs. The “all accumulators” network received an additional input for each evidence accumulator, conveying its value (Fig. 2). The “max accumulator” only received one additional input, the value of the maximum accumulator. This network is related to the race model of stopping, according to which animals use a stopping criterion on the value of the maximum accumulator (Bogacz et al., 2006). The “all probabilities” network received an additional input for each choice, in addition to time. These inputs corresponded to the posterior probability of each choice. The “max probability” network only received one additional input, the value of maximum posterior probability. Hence, this network received only the variables which are needed for the approximately optimal stopping criterion.

In each training epoch we presented the networks with the input data for every trial, and every time step. The activation of the output unit in response to the input data for trial *i* and time step *t* was interpreted as the probability that the network stopped and decided at time step *t* in trial *i*, if the network had not already stopped and decided. The tanh activation of the output unit lead to activations in the range (−1, 1), so we mapped linearly to the range (0, 1) to provide a number which could be interpreted as a probability. On the final time step, step 40, the networks were forced to stop and decide.

The networks were trained to minimise the average loss across all trials and time steps. The loss is the negative reward earned by the network. Once the network “stopped” we assumed the network then made the Bayes optimal decision, and the network received the corresponding reward. From this reward we subtracted the cost of time used to accumulate until the time step of the decision. As we interpreted the output of the network probabilistically, we did not enforce that the network stop at a specific time. Instead, our interpretation of the output gives us a probability distribution over stopping times. The network earned the reward associated with a particular stopping time in proportion to the probability mass assigned by the network to that stopping time. Hence, the reward earned by the network represents the expected reward under a probabilistic stopping rule.

We prepared the training data for the constant cost of time condition by *z*-scoring the evidence values for all trials, all time steps and all choices together, and separately *z*-scoring the time step value for all trials and all time steps together (LeCun, Bottou, Orr & Müller, 1998). The cost of time does not affect the distribution from which training data is drawn. In preparing data for the increasing cost of time condition, we did not *z*-score but applied the same centring and scaling values used for the constant cost of time condition. (Each centring and scaling from the constant cost of time condition was applied once in the increasing condition.)

When assessing the performance of the networks we used the reward obtained per trial in the validation sets (the part of the simulated data sets not used for training). All plots show this reward. In order to obtain an estimate of the uncertainty associated with the performance of the networks, we trained each network 20 times (from random initial weights).

### Hyperparameter optimisation

An issue with using neural networks as model learners is that they introduce an element of arbitrariness. One must select the parameters of the networks, and the concern is that this may lead to conclusions which depend on those parameters. To alleviate this, we used the Hyperopt Python library to perform hyperparameter optimisation (Bergstra, Yamins & Cox, 2013). Hyperparameter optimisation involves scanning over the network hyperparameters and choosing the configuration which maximises some objective function. Thus the choice of network properties is, to some extent, taken out of the hands of the experimenter.

We optimised over the parameters that plausibly have a large impact on the results; for example, a high learning rate may impair a network’s ability to find the global minimum, whilst a very low learning rate may artificially slow down a network. This becomes a particular problem if different networks perform best at different learning rates. By optimising over learning rate, we can reduce the probability that there are such effects in our results. For the hyperparameter optimisation options considered, see Table 2. Every possible combination of parameters was considered at least once.

**Table 2:** Network parameters used for the model learner. For parameters which were optimised, we indicate the possible choices that were scanned over.

| Network variable | Parameter used | Hyperparameter optimisation choices |
| --- | --- | --- |
| Optimiser | RMSprop | Adam, RMSprop, SGD, Adadelta |
| Learning rate | 0.0005 | 0.0001, 0.0005, 0.001 |
| Dropout rate | 0.5 | 0.2, 0.5 |
| Number of layers | 4 | 3, 4, 5, 6, 7, 8, 11 |
| Network type | Dense, fully connected | - |
| Training-validation split | 75:25 | - |
| Activation function | tanh | - |

Our objective function was the reward obtained in 3000 unseen trials, after 500 epochs of training on data from the 4 choice condition, with a constant cost of time, and no correlation between accumulators. Using previously unseen trials allows us to avoid parameter combinations which produce overfitting. Importantly, we performed hyperparameter optimisation using the “all accumulators” network. That is, network properties were selected as those which allowed this network to perform best. Hence, if any network had an unfair advantage due to the specific network settings used, it was this one, and any increased performance we see in our favoured model, “max probability”, is a conservative estimate of the improved performance compared to “all accumulators”. Hyperparameter optimisation produced the choice of network parameters shown in Table 2.

### Code availability

Upon publication, all code written for the study will be made available at a doi to be determined.

## 4 Modelling results

We trained neural networks on a simulated perceptual decision making task, in which the networks needed to learn a policy for stopping evidence accumulation and triggering a decision. Here we discuss the results for the the simplest (2-way choice) and most difficult (10-way choice) cases. Results were similar for the intermediate difficulty (4-way choice), and we provide these in supplementary file S3.

### 4.1 2 options

After training the networks we explored the central question of whether learning the mapping from time and state of the evidence accumulators, to stopping vs. waiting, can be sped up through approximation. Figs. 3 shows the time course, over rounds (epochs) of training, of the performance of the networks. The figure shows this time course in environments with increasing and constant cost functions, and with uncorrelated and correlated evidence accumulators. Lines represent the median reward obtained by the network over 20 attempts to train the network (from random weights), and shading represents 95% confidence intervals computed using bootstrapping.

**Figure 3:**
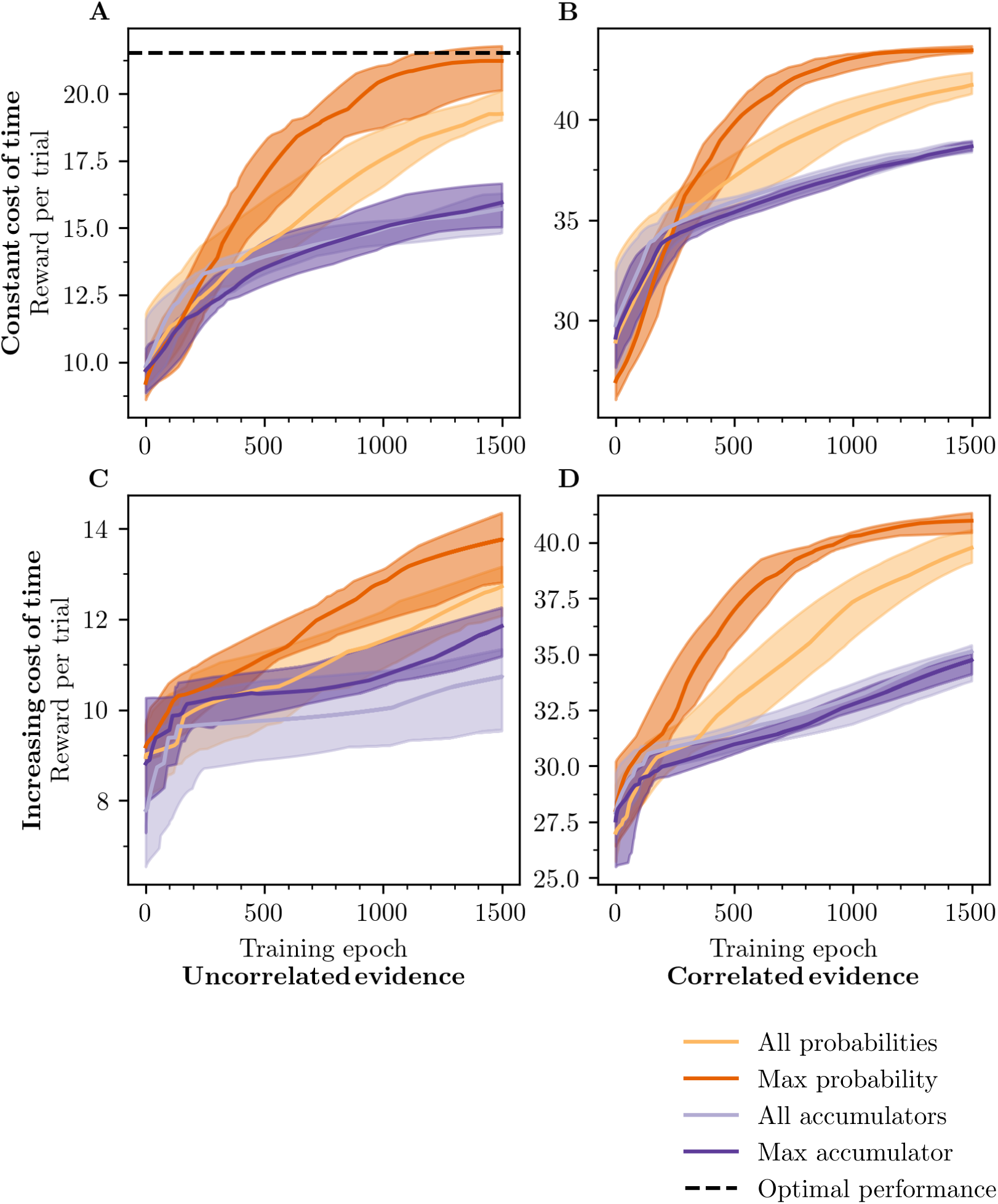
Average reward gained per trial for the case of 2 choices. Results are shown for cases with and without correlations between evidence accumulations, and with a constant and an increasing cost of time. The optimal performance for the uncorrelated constant cost of time case is calculated according to the discussion in supplementary file S2.

Under no condition tested was the “max probability” network outperformed; in all cases it learned as fast or faster than all other networks. In the case of uncorrelated evidence with a constant cost of time (Fig. 3, A), it is the “max probability” network which is able to achieve the optimal reward within 1500 training epochs. These results suggest that approximating the optimal stopping criterion on time and the state of all accumulators with a stopping criterion on time and the maximum posterior probability, leads to faster learning.

Interestingly, in most cases, the “all probabilities” network learned faster than the “all accumulators” network, suggesting that using probabilities enables faster learning than when using the raw accumulated evidence samples. The dimensionality of input for both networks is the same, although the probabilities are constrained to a line such that *p*(*C* = 1|***x***) + *p*(*C* = 2|***x***) = 1. The raw evidence samples are not so constrained, despite the fact that the sum of evidence samples, ∑_*i*_ *x*_*i*_(*t*), is irrelevant to the question of when to stop (Tajima et al., 2019). (This sum is irrelevant because adding the same amount of extra evidence to all accumulators does not change the posterior probabilities, and because of this it does not change the future behaviour of the accumulators relative to their current point, nor the value of deciding at those future points; see equations 35, 36, 9.) This additional unnecessary variability may explain why it is harder to learn a mapping from the raw evidence samples, than from probabilities.

We note that the approximation embodied by the “max probability” network does not lead to fast learning simply because the approximation reduces the dimensionality of the input for the mapping that must be learned. The “max accumulator” network also learns a mapping from two dimensions (time and the maximum accumulator) but this network learns far slower than even the “all probabilities” network, which learns a mapping from a three dimensional space.

One may wonder why the networks achieve a greater reward in the correlated environment, rather than in the uncorrelated environment. This effect arises because, when the evidence accumulators are correlated, a greater proportion of variability is along the direction [1, 1]^*T*^. The value of the state of the accumulators along this dimension gives the value of the sum of the accumulators and, as discussed, this sum is irrelevant to the posterior probabilities for the two options. Hence, in the correlated case a larger proportion of variability is along a dimension which does not affect how discriminable the two options are, making the task easier.

Taken together these results suggest that animals, if they are aiming to rapidly learn optimal stopping, should learn a mapping from time and maximum posterior probability. Using the full posterior, while it offers no dimension reduction is also a sensible strategy when faced with 2-choices. In either case, animals should use the products of Bayesian inference in their criteria for stopping.

We additionally explored the decision rule learned by each of the networks. To do this we simulated new trials, and looked at how the networks behaved in these new trials. For each trial we found the networks’ probabilistic policy by interpreting the output in the usual way (see Sec. 3), and simulated the behaviour of an observer which followed this policy. Hence, for each trial we found one stopping time, drawn according to the probabilistic policy. For each trial, and each time step at which the simulated agent waited, we plotted the difference in evidence for the two choices (accumulator difference), against time. We also plotted these values at the time step when the agent made a decision. We show the results of this analysis in Fig. 4, for the “max probability” and “all accumulator” networks, in the case where there is correlation between accumulators.

**Figure 4:**
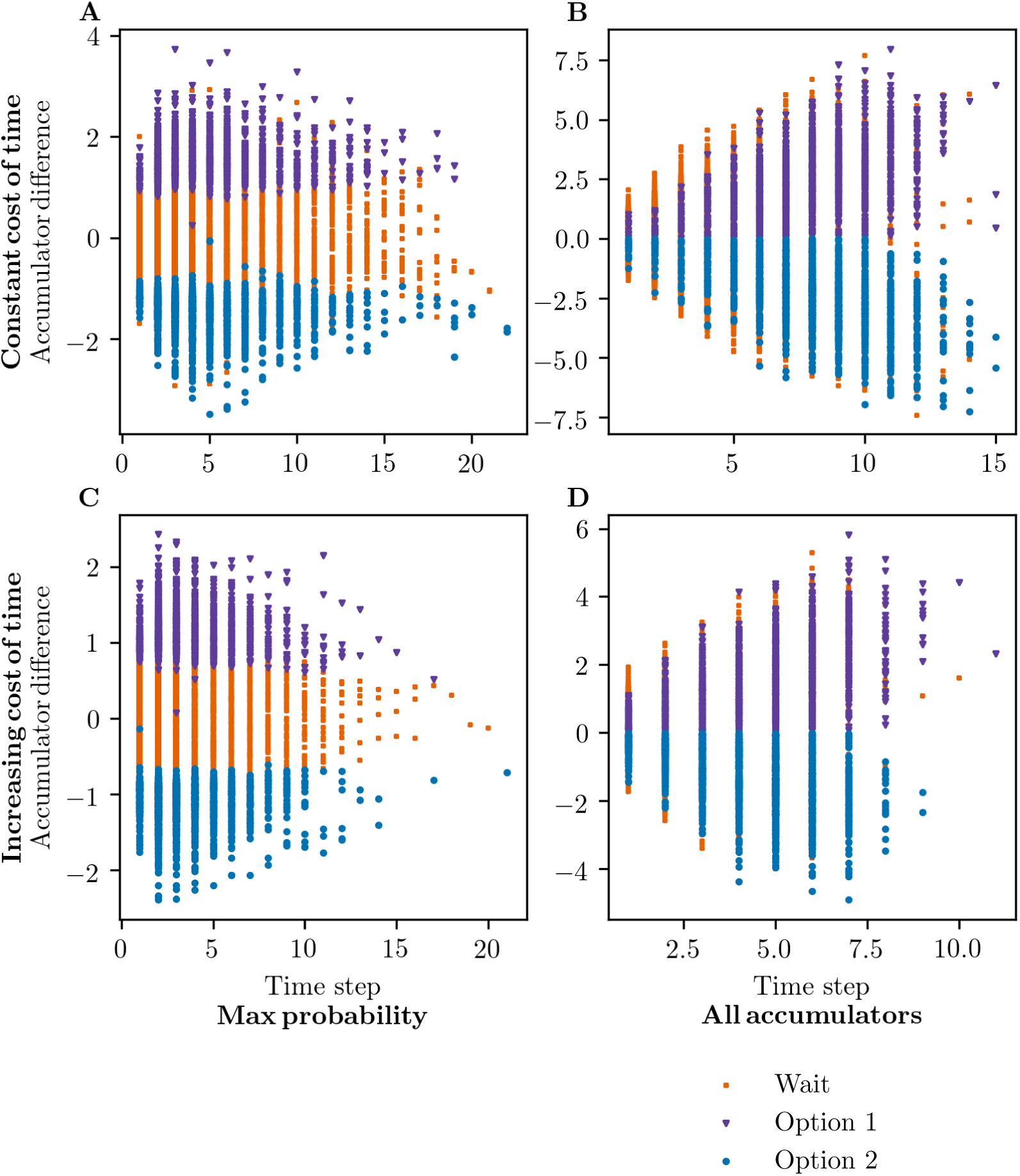
Learned decision rule for “max probability” (left) and “all accumulators” (right) networks, for correlated evidence accumulations with a constant cost of time (top) or increasing cost of time (bottom) for the case of 2 choices. Accumulator difference refers to the difference in the values of the two evidence accumulators.

When the cost of time is constant, the optimal policy does not depend on time (Ferguson, 2011). This is because, whether you are at time step 5 or time step 10, if the current posterior is the same, the probability distribution over the evolution of the accumulator state in future time steps, and the cost of time in future time steps is identical. Certainly at time step 10 you will already have incurred a greater cost of waiting than at time step 5, but these are sunk costs you will have to pay whether you wait or stop, and hence, do not affect whether waiting or stopping will provide greater reward. In the case we consider, the difference between accumulators is monotonically related to the posterior probabilities (see equation 35; Moran, 2015). Therefore, the networks should require the same difference between accumulators before they stop, regardless of how long they have been deliberating.

For the “max probability” model, we observe that, as expected, the difference in evidence required to make a decision is approximately constant with time (Fig. 4; A). It is interesting to note that this is the policy used in the prominent diffusion decision model (Ratcliff & McKoon, 2008). The “all accumulator” network learned a very different rule, reflecting that fact that even after 1500 training epochs, this network was still performing well below the level of other networks (Fig. 4; B). The network had not learned a clear policy, and the network stopped accumulation and made decisions even when the difference in evidence between the two alternatives was zero; i.e. it was deciding despite having no evidence to favour either option.

Under conditions in which the cost of time is increasing, we expect the networks to require less evidence as time increases, as it becomes increasingly detrimental for the network to wait. Again the “all accumulator” network did not learn a clear policy (Fig. 4; D). For the “max probability” network it is difficult to see from the plot whether the correct stopping rule, with a decreasing criterion on the difference in evidence, has been learned (Fig. 4; C). The policy learned by the network becomes clearer if we plot it in a different way. Instead of plotting particular simulated behaviours, we plot the probability the network stops, as a function of the difference between the two accumulators, and the time step (Fig. 5). From these plots it is clear that the “max probability” network has learned a stopping policy which features a decreasing stopping criterion on the difference in accumulated evidence (Fig. 5; B). It is interesting to note that the network even learns a sensible policy for time steps which it rarely encounters (compare with the time steps reached in Fig. 4; C).

**Figure 5:**
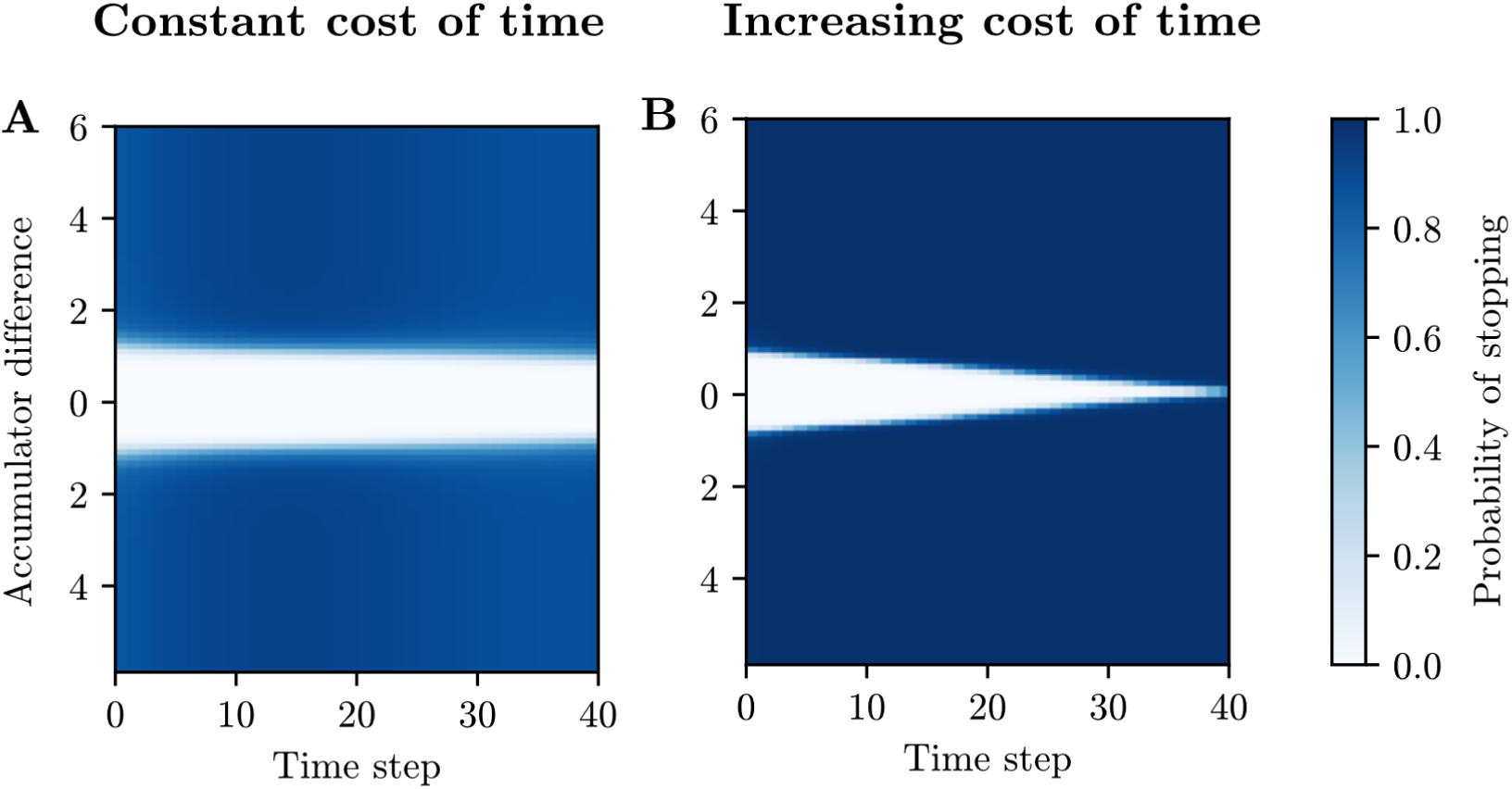
Probability the “max probability” network stops as a function of the difference between evidence accumulators, and time. Results shown for 2 choice options, correlated evidence accumulators and with a constant (A) or increasing (B) cost of time.

These results suggest that, in the case of two choices, learning an approximation to the full mapping from time and state of the accumulators, to stopping vs. waiting, can speed up learning. Learning the mapping from time and maximum posterior probability is particularly fast. Furthermore, learning such an approximate mapping leads to a sensible stopping policy, suited to the current conditions.

### 4.2 10 options

The “max probability” network outperforms the other networks when choosing between 2 options. We also considered the much more challenging situation in which the network had to choose between 10 options. In Fig. 6 we plot the average reward per trial achieved by the different networks, over the course of training. Again the “max probability” network learned as fast, or faster than the other networks. In some cases the advantage was very large (Fig. 6, D). The only case in which the “max probability” network did not learn faster was with uncorrelated evidence samples and an increasing cost of time. As discussed in the previous section, the task with uncorrelated evidence samples is considerably harder than the task with correlated evidence samples. Coupled with the increasing cost of time, it may simply be that there is little for any network to learn in this case.

**Figure 6:**
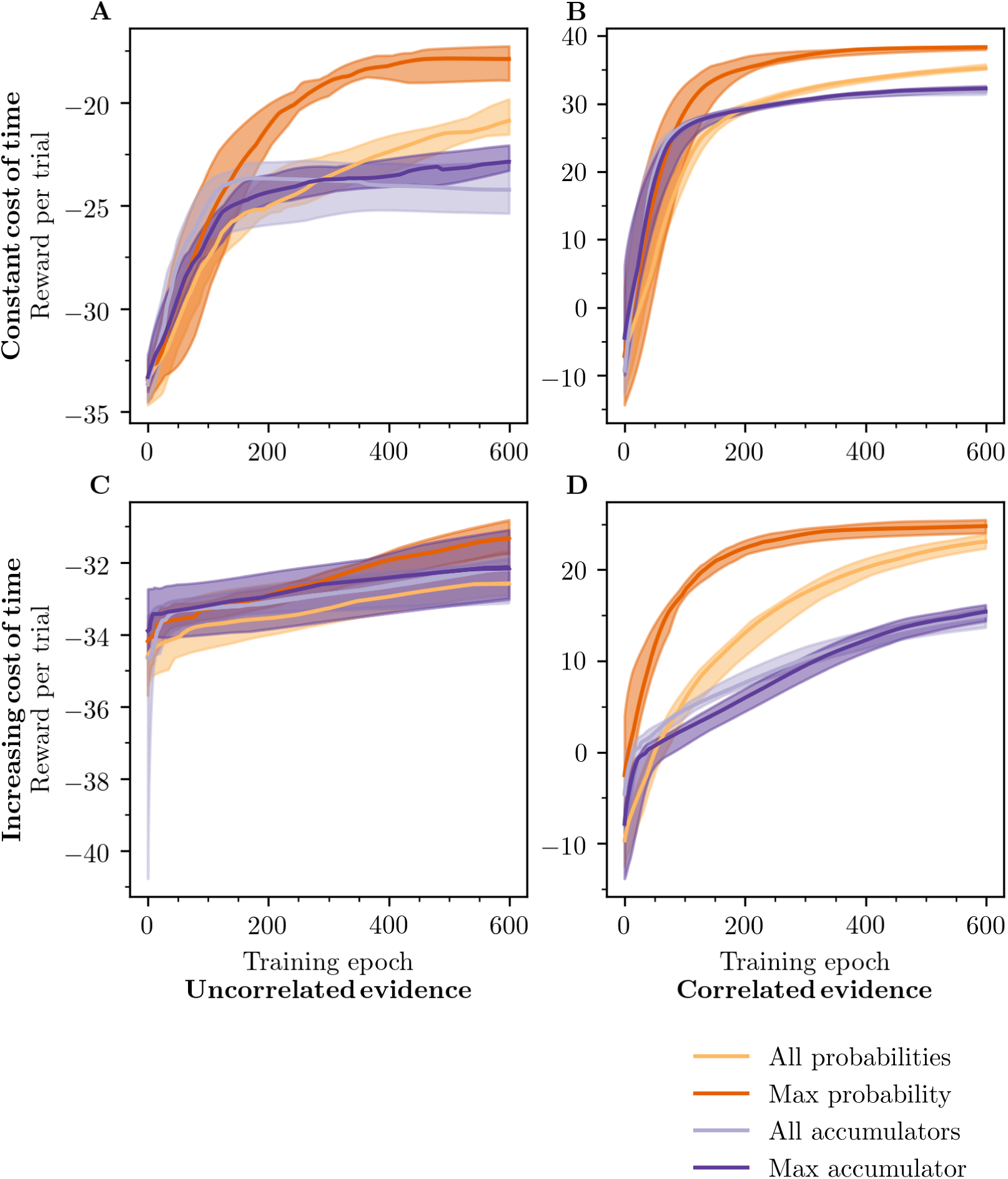
Average reward gained per trial for the case of 10 choices. Results are shown for cases with and without correlations between evidence accumulators, and with a constant and an increasing cost of time.

With 10 accumulators, one concern might be that the benefit of the approximation embodied by the “max probability” model is largely down to the fact that this approximation reduces the dimensionality of the input space from 11 (time and the 10 accumulators) to 2. However, we again see that the “max accumulator” network, which also learns a mapping from 2 dimensions (time and the maximum accumulator), performs far worse than the “max probability” network. Hence, the specific information represented by the maximum posterior probability seems to be the key to rapid learning.

As before, we are also able to check whether the networks are able to learn a sensible stopping policy. It is more difficult to visualise the learned behaviour with 10 choices because of the dimensionality of the evidence space. We have to reduce the dimensionality of the cumulative evidence (10 values) in order to visualise the decision rule. We pick two options at random and plot the behaviour against the difference in the evidence accumulators for these two options. Naturally a lot of information is lost by doing this.

In Fig. 7 we show plots of the difference between evidence accumulators for two randomly selected accumulators, and behaviour at that difference, against time, as we did in the previous section. The “max probability” network learns to behave sensibly; if there is no evidence to distinguish between the two options, the network does not pick one of them (Fig. 7, A). In contrast, the “all accumulators” network stops and decides even when there is no difference in evidence between the chosen option and the other option used for plotting (Fig. 7, B, D). This might be a sensible thing to do if the cost of time is increasing, however, the network also behaves this way when the cost of time is constant. When the cost of time is constant, the network should use the same stopping rule at all time steps, as discussed. However, the “all accumulators” network clearly violates this feature of the optimal solution (Fig. 7, B).

**Figure 7:**
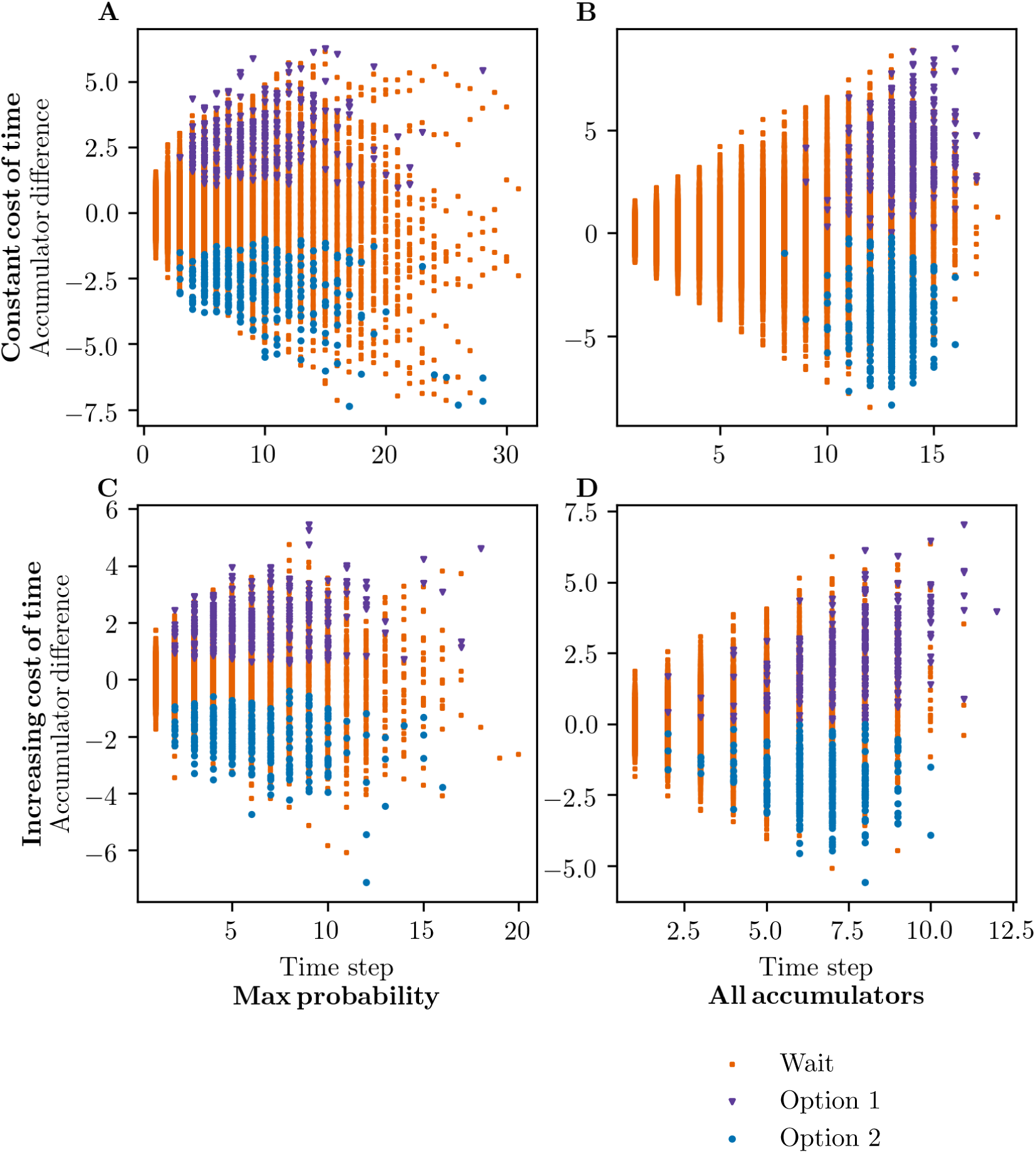
Learned decision rule for “max probability” (left) and “all accumulators” (right) networks, for correlated evidence accumulators with a constant cost of time (top) or increasing cost of time (bottom) for the case of 10 choices.

In conjunction with the results from the 2 options case, these findings suggest that learning an approximate stopping rule by using a criterion on time and maximum posterior probability, leads both to sensible stopping policies, and considerably faster learning.

## 5 Discussion

We considered the problem of optimal stopping in a generic perceptual decision making task. We found that under a certain approximation – considering only the most probable events at future time steps – only time and the maximum posterior probability are relevant to the question of when to stop deliberation.

We explored whether the simplification afforded by using a criterion on only two variables (time and maximum posterior probability) sped up learning. We compared several networks, including a network which learned a stopping criterion on time and the full description of the accumulation state (the value of all accumulators). The network which used a criterion on only time and maximum posterior probability, and hence used the structure of the approximate optimal stopping rule, learned fastest. We found evidence that it is the maximum posterior probability specifically, and not simply the dimensionality reduction, which supports the rapid learning of sensible and near optimal stopping rules. Specifically, a network which learned a mapping from time and the maximum accumulator performed poorly.

These results provide an interesting link between approximately optimal stopping, and optimal deciding. Intuitively, an animal’s probabilistic estimate of the state of the world should not just be relevant to the question of what to decide, but also when to decide. In the context of decisions between more than two alternatives, only deciding has been extensively studied using the idea that animals perform Bayesian inference (although there are exceptions e.g. Bogacz and Gurney, 2007). Our results suggest that variables computed during Bayesian inference are also important for approximately optimal stopping.

This overlap of the variables needed for optimal stopping and optimal deciding provides a clear opportunity for animals with limited neural processing capacity; the same neural populations which are used in optimal deciding can be used be used to support optimal stopping. A corollary of this conclusion is that, if we believe animals are computing posterior distributions for the purpose of deciding, it seems unlikely that they are not recycling these distributions when such re-use could provide great benefit for learning to stop.

Conceptually, our work is largely consistent and inspired by the work of Tajima et al. (2019), who showed that animals could approximate the optimal stopping rule by mapping the state of the accumulators onto a manifold. Our contribution is, first, to consider how approximation might affect the speed of acquisition, and second, to tie optimal stopping to Bayesian inference, by showing how conversion of accumulators to posterior probability is a specific mapping which can both be theoretically motivated, and conveys practical benefits to animals.

The idea that animals might use stopping criteria involving posterior probabilities has been proposed before. Stopping rules involving probabilities have been studied as sensible approaches to statistical problems (Baum & Veeravalli, 1994; Dragalin et al., 1999). Bogacz and Gurney (2007) took one of these rules, a form of “multihypothesis sequential probability ratio test” (MSPRT), and considered how it might be implemented in the brain. Using this rule, animals would continue accumulating evidence until the maximum posterior probability reaches a threshold which is constant over time. This policy is only optimal in specific situations (Dragalin et al., 1999). In general, a time-dependent threshold is required for optimal stopping (Tajima et al., 2019). Our work builds on the idea that the brain implements a MSPRT, by suggesting there are time-dependent thresholds on the maximum posterior probability, and by showing that these provide approximately optimal stopping in a wide range of conditions.

While we didn’t directly consider the dominant model of stopping, the diffusion decision model (DDM; Ratcliff and McKoon, 2008), which only applies in two-alternative decision making tasks, we note that the principles of the DDM are very similar to the principles embodied by one of the networks studied. In the DDM, a response is triggered when the difference in evidence accumulated for the two alternatives reaches a threshold. At any particular time step, the posterior ratio between the two alternatives is monotonically related to the difference in evidence accumulated for the two options (Bogacz et al., 2006; Moran, 2015). Hence, in the DDM, observers apply a stopping criterion on time and the posterior probabilities. In the DDM the decision boundaries do not necessarily need to be symmetric (Ratcliff & McKoon, 2008), hence at any particular time step decisions for the two options might not be made with the same posterior probability. We studied a network which learned a criterion on time, and the posterior probabilities of the alternatives. As this network received information on which option had the highest posterior probability, it could learn asymmetric decision boundaries. We found this network performed reasonably, especially in the case of two choices, and outperformed networks which learned criteria on the accumulator states.

The race model is also related to one of the networks used in this study. The race model of stopping suggests that it is the maximum accumulator which triggers a response (Bogacz et al., 2006). We tested a network which used time and the maximum accumulator in its stopping rule, however this network learned far more slowly than networks which used probabilities in their stopping rules. These results suggest that an animal which used a time-dependent race model would suffer from slower learning, and hence, a decrease in efficiency and/or quality of decisions.

Taking a wider perspective, our work adds to findings suggesting that decision confidence is a variable of very general importance. Confidence has been found to be important in a range of processes such as information-seeking (Desender, Boldt & Yeung, 2018), group decision making (Bahrami et al., 2010), sequences of related decisions (van den Berg, Zylberberg, Kiani, Shadlen & Wolpert, 2016), and learning useful low-dimensional representations of stimuli (Drugowitsch, Mendonça, Mainen & Pouget, 2018). We found that animals can learn an approximately optimal stopping rule by using a criterion on time and maximum posterior probability. The maximum posterior probability is the probability of being correct if a decision is made, and the quantity which optimally calibrated confidence reflects (Kepecs & Mainen, 2012). Hence, our work provides another process where a representation of decision confidence may be important.

The study has two main limitations. First, the conclusions drawn only directly apply to perceptual decisions of the form studied. We considered a decision in which the observer could have any prior belief about the options, we allowed correlations between the accumulators, and allowed any function for how the cost of continued deliberation changes with time. When creating any model, a large number of assumptions will have to be made. Perhaps the most important ones in our case are that we did not consider arbitrary covariance structures, different rewards associated with different options, or variability in the signal-to-noise ratio (*µ*). We anticipate that in these cases similar principles to those identified here will apply; with a sensible approximation it will be possible to reduce high dimensional problems down to a few dimensions, and posterior probabilities over the state of the world will have a key role to play in this.

The second main limitation is the nature of the neural networks. We worked to ensure maximum generalisability of our neural network results by selecting neural network properties as those which most helped a competitor to our favoured network (see “Hyperparameter optimisation” above). In any case, there are features of the network training which are unlike the process animals and humans engage in. Specifically, the networks did not have to commit to a single decision, but provided a probability of stopping at each time step, and received feedback on this probabilistic policy. Hence, the networks received more information about good and bad behaviours than an animal that stops at a single time step, and can only learn about the value of stopping at this one step. We were not aiming to provide realistic models of the mechanism by which animals and humans learn. Instead we were aiming to test the idea that approximation could speed up learning in general. In fact, if learning in animals and humans is more protracted than learning in the networks used here, the importance of using the approximate solution to speed up learning is likely to be even greater than it appears here.

Our findings provide evidence that the products of Bayesian inference may be even more useful than previously thought. Using the maximum posterior probability, animals may be able to simplify stopping problems and speed up learning, providing evolutionary advantages, especially in changing environments.

## Acknowledgements

ES has received funding from the European Research Council (ERC) under the European Union’s Horizon 2020 research and innovation programme (grant agreement No 788223, PanScales) and from the ERC Starting Grant PDF4BSM. JCT would like to thank the members of the ACC lab (Oxford) for helpful comments on the work.

## Competing interests

The authors have no competing interests.

## S1 Mathematical results

### S1.1 Decision problem and posterior

As discussed in the main text, every time step the observer recieves evidence samples, *δ****x***(*t*), according to

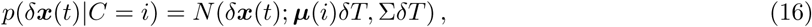

where *t* indicates the current time step, *δT* is the duration of a time step, *N* () indicates the multivariate normal distribution, and Σ is the covariance of evidence samples (Tajima et al., 2019).

We want to compute the posterior over *C* using Bayes rule. Before we can use Bayes rule we need to know the likelihood function of *C*, given evidence samples. After *M* time steps the observer will have gathered evidence samples *δ****x***(1), *δ****x***(2), …, *δ****x***(*M*). Using the independence between the samples gathered at different time steps (conditioned on the value of *C*), the likelihood will be

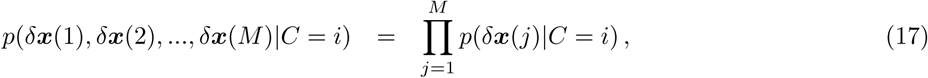

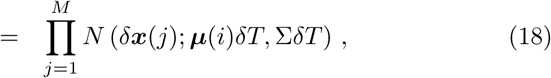

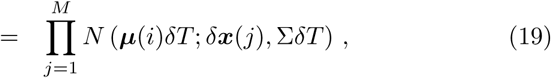

where we have used (16).

Using standard results for the product of multivariate normal distributions,

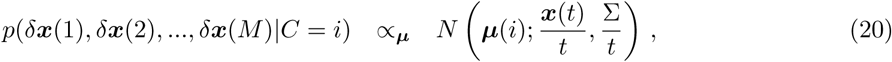

where ***x***(*t*) =∑_*i*_ *δ****x***(*i*), *t* = ∑_*i*_ *δT* and the proportionality is with respect to *µ* (Drugowitsch et al., 2012).

We can now calculate the posterior

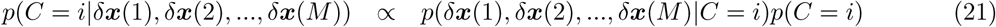

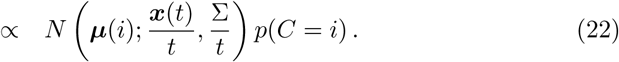

Note that

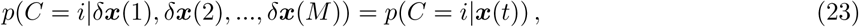

consistent with the conclusions of Moreno-Bote (2010). We find a different form for the posterior by considering the log-posterior ratio

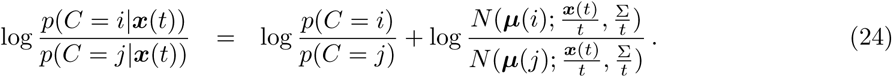

We consider only covariance matrices which can be written in the following form

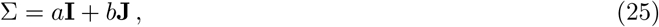

where **J** is a matrix of ones. We can write this covariance matrix as

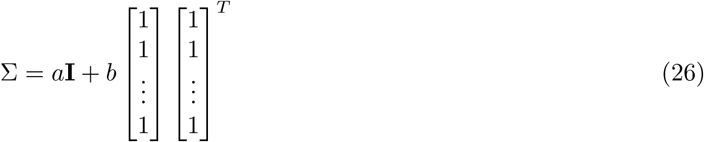

and use the Sherman–Morrison formula to invert it, giving

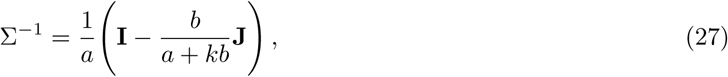

where *k* is the dimensionality of ***x*** (and therefore, the number of possible values of *C*).

This allows us to rewrite the normal distribution

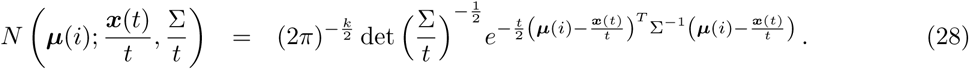

Consider the exponent

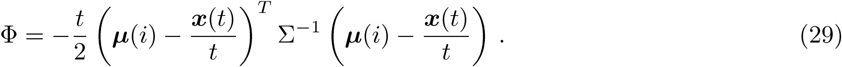

Using (27), we have

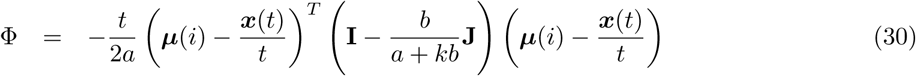

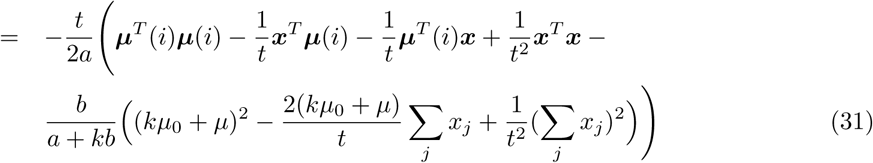

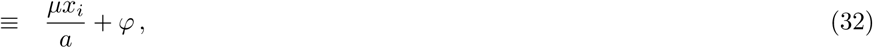

where *x*_*i*_ is the *i*th element of ***x***, and *φ* includes all terms which do not depend on *i*.

We can use this result in our expression for the log-posterior ratio (24),

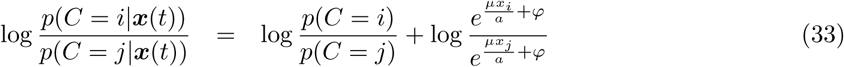

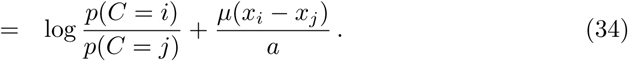

Using the fact that Σ_*i*_ *p*(*C* = *i*|***x***(*t*)) = 1, we have

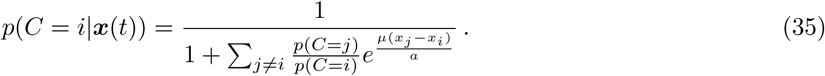

### S1.2 Approximating the optimal solution

The future evolution of ***x***(*t*) is unknown, but described by

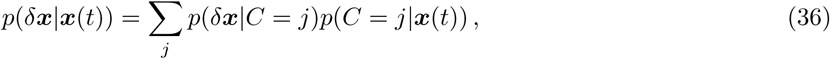

where *δ****x*** is the change in ***x*** at any future time step. We approximate the future probability distribution over ***x*** by, at each time step, only considering the most likely value for *δ****x***. This reduces the problem of picking a stopping policy to the problem of picking a time to stop.

The *p*(*δ****x***|*C* = *j*) terms in (36) are multivariate normal distributions given by (16). The maximum value reached by each is the same, because the covariance of these multivariate distributions does not depend on *j*. Hence, the most probable value for *δ****x*** is determined by the multivariate normal distribution which is multiplied by the *p*(*C* = *j*|***x***(*t*)) term which is greatest. The maximum value achieved by the corresponding multivariate normal distribution will be at its mean, ***µ***(*j*)*δT*. That is,

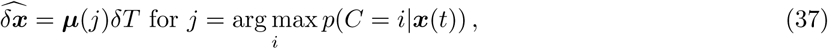

where 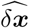 is the most likely value of *δ****x***.

Note that the most likely value of *δ****x*** is such that *δ****x*** will only provide additional evidence for the most probable option. Hence, if *C* = *j* is currently the most probable option, under our approximation, it always will be.

Consider two points ***x***^(*d*)^(*t*) and ***x***^(*e*)^(*t*) for which the posterior probability of the most probable option is the same. That is,

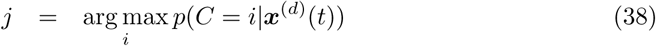

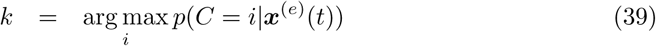

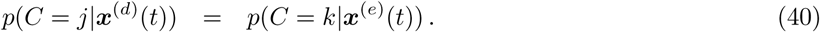

Using (35) we have

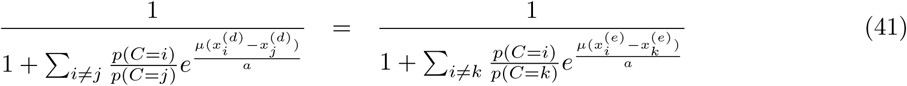

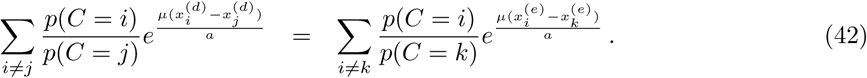

Consider some later time step *t* + *l*. Under our approximation,

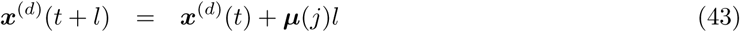

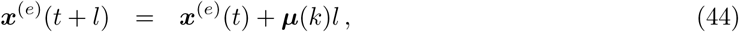

and hence considering components of the vector ***x***,

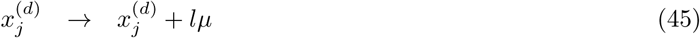

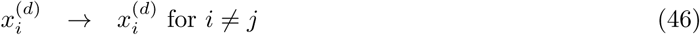

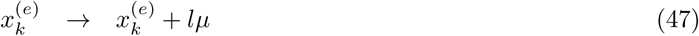

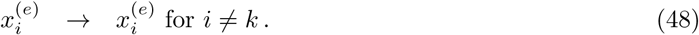

Under the approximation, the expected reward for starting from ***x***^(*d*)^(*t*) and stopping at *t* + *l* is, recalling (9), given by

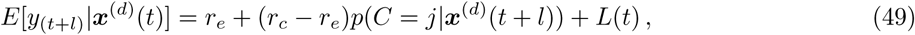

because *C* = *j* is the most probable option at time step *t*, and we know that under our approximation, the most likely option will not change. Again using, (35), and using our expressions for the components of ***x***(*t* + *l*),

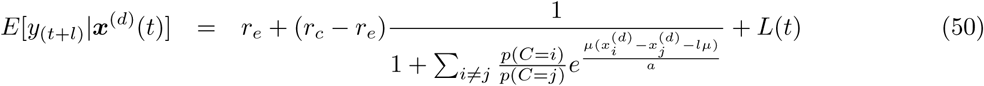

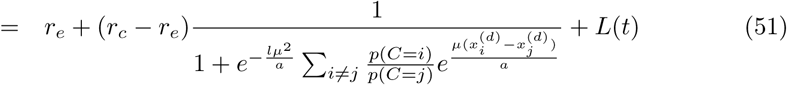

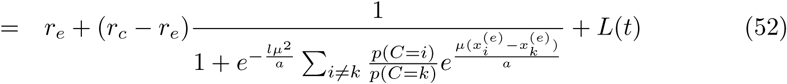

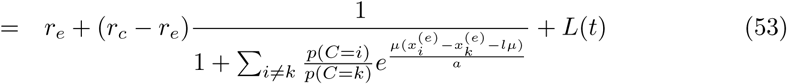

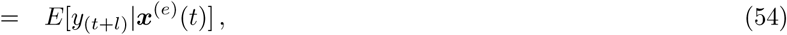

where we have used equation (42) moving from line two to three. This result shows that if the maximum posterior probability at two points is equal at time step *t*, then the expected reward at any later time step *t* + *l* is the same under our approximation.

## S2 Optimal rule for simple case

In the case of independent accumulators, 2 response options, and a constant cost of time, we can calculate performance under the optimal stopping rule.

As discussed in the main text (Sec. 4.1), when the cost of time is constant, the optimal stopping rule, when described as a criterion on the posterior probabilities, does not change with time (Ferguson, 2011). In our case, the posterior probabilities are monotonically related to the difference between the two accumulators (see equation (35)). Hence, the problem of finding the optimal stopping rule reduces to the problem of finding the height of a time-independent stopping threshold on the difference between the accumulators.

Denote *x* = *x*_1_ − *x*_2_, where *x*_*i*_ represents the *i*th element of ***x***, which is defined in the main text. We are looking for the optimal magnitude of *x* at which to stop and decide. That is, we will only make choices for *C* = 1 at *x* = *K*, and choices for *C* = 2 at *x* = −*K*. Using equation (35), an equal prior, and substituting in *x* we have

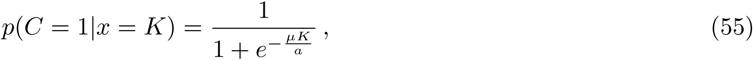

and a similar result holds for *C* = 2. If we are in a trial in which we reach *x* = *K* and hence the response is *Ĉ* = 1, then the probability we are correct is *p*(*C* = 1|*x* = *K*). In the perceptual decision on which the networks were trained, the reward for getting the wrong answer was the negative of the reward for getting the right answer. If we denote these rewards −*r*, and *r*, if we therefore reach *x* = *K* the expected reward from deciding is

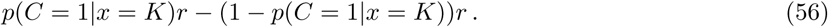

A similar result holds for a response *Ĉ* = 2 at the boundary *x* = −*K*, and by symmetry *p*(*C* = 1|*x* = *K*) = *p*(*C* = 2|*x* = −*K*). Denote this value *P* = *p*(*C* = 1|*x* = *K*) = *p*(*C* = 2|*x* = −*K*), and note that *P* is the probability of making the correct decision. Hence, regardless of the boundary reached, the expected reward from deciding is

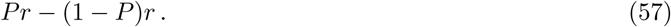

In order to calculate the overall expected reward, we also need to know the cost of time, and hence the expected number of time steps spent accumulating evidence. From Shadlen, Hanks, Churchland, Kiani and Yang (2006) we have that the expected number of time steps equals the expected value of *x* at decision time, divided by the average change in *x* in the absence of decision boundaries. When the true value of *C* is *C* = 1, the average change in *x* = *x*_1_ − *x*_2_ in the absence of boundaries is just *µ*. Hence,

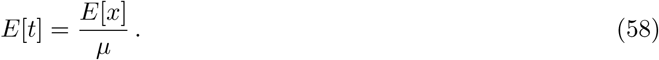

When *C* = 1, the correct decision boundary is *x* = *K* and the probability of reaching this boundary is *P*. There is a 1 − *P* probability of reaching the other boundary at *x* = −*K*. Hence,

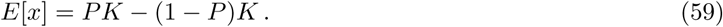

Which gives us

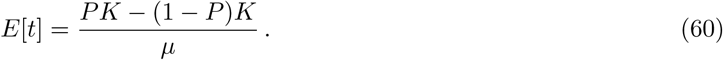

An identical expression holds when *C* = 2.

The overall expected reward, *E*[*y*], is the expected reward from deciding minus the expected cost of time

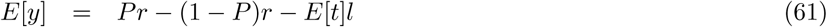

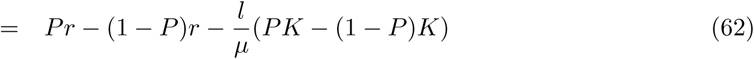

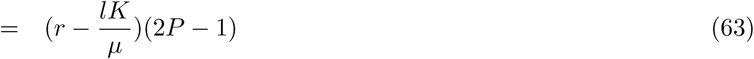

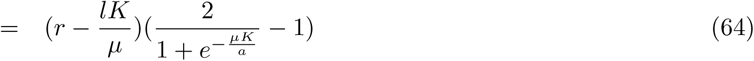

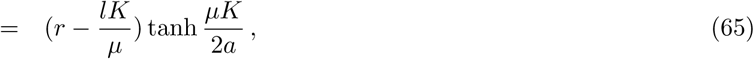

where *l* is the cost per time step.

To find the value of K which maximises the expected reward we take the derivative. Setting the derivative to zero and rearranging we find

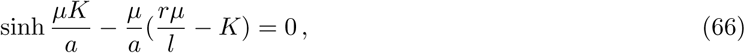

which we can solve for *K* using standard solvers. Using *K* we can find the maximum obtainable reward using equation (65).

## S3 4 options

In addition to requiring the neural networks to choose between 2 and 10 options, we also looked at an intermediate case, where the networks had to choose between 4 options. In Fig. 8 we again show the reward per trial achieved by the different networks. Unsurprisingly, the behaviour of the networks is somewhere between the behaviour observed for 2 and 10 locations. Again, in all cases, the “all probability” network is as good as, or better than all other networks; for all cases it is able to learn the optimal stopping rule the fastest.

**Figure 8:**
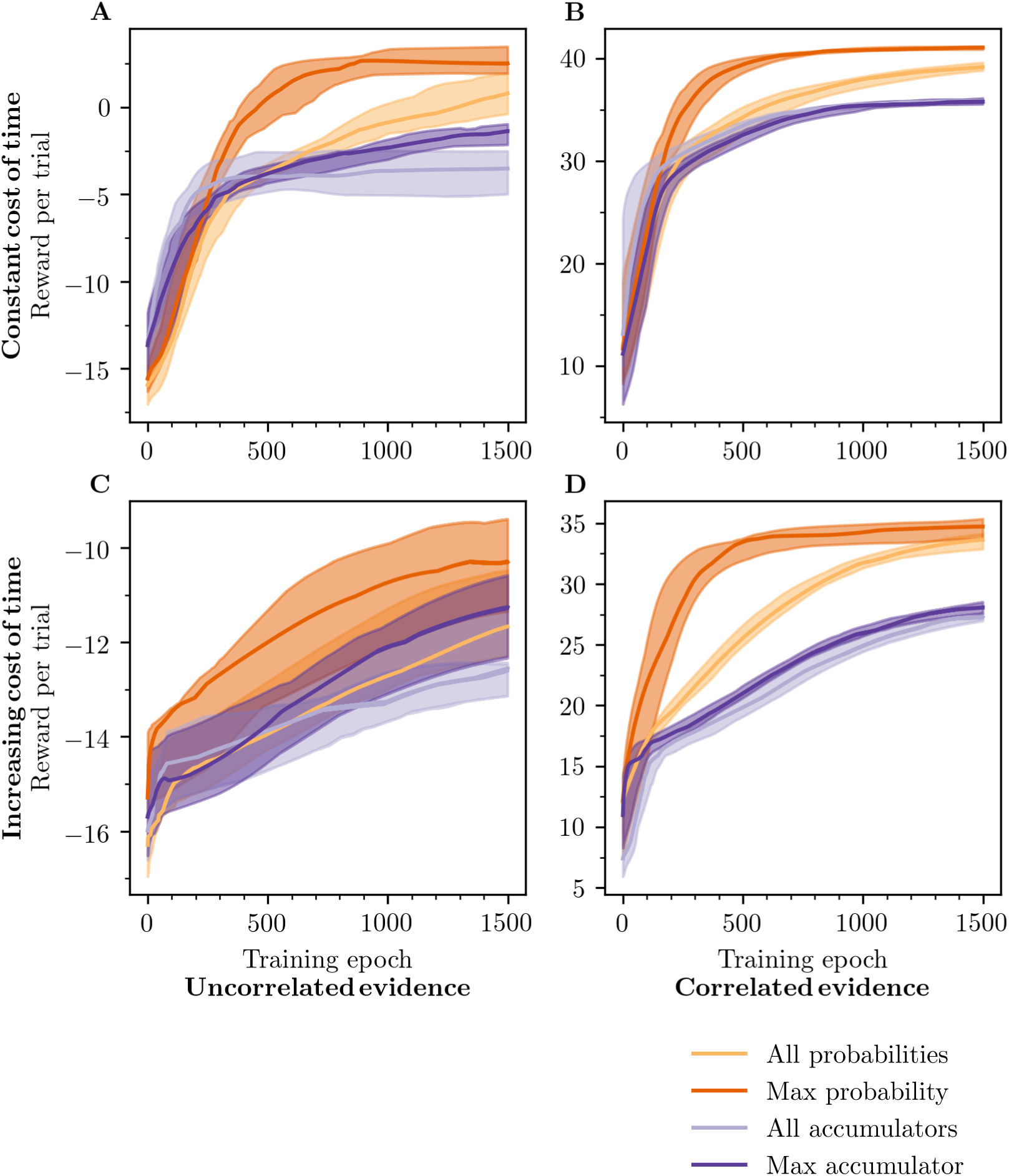
Average reward gained per trial for the case of 4 choices. Results are shown for cases with and without correlations between evidence accumulations, and with a constant and an increasing cost of time.

In Fig. 9 we show show the learned stopping rules for the “max probability” and “all accumulator” networks. In the case of a constant cost of time (Fig. 9, A, B), the “all accumulator” model is again not learning a sensible stopping rule, whilst the “max probability” network has learned a rule which is consistent with the features of optimal stopping discussed in the main text. There is a similar pattern of results in the increasing cost case, (Fig. 9, C, D).

**Figure 9:**
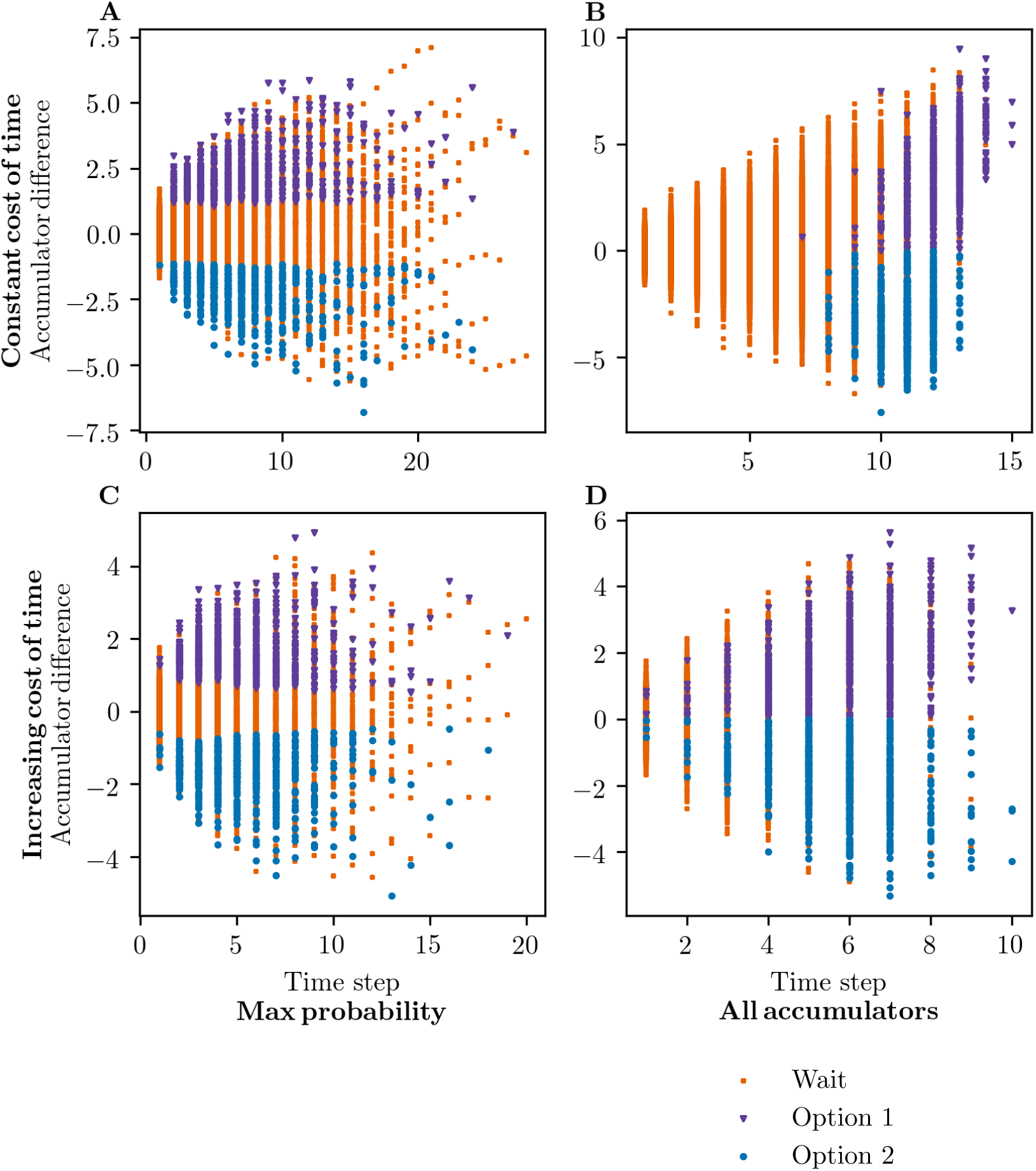
Learned decision rule for “max probability” (left) and “all accumulator” (right) networks, for correlated evidence accumulations with a constant cost of time (top) or increasing cost of time (bottom) for the case of 4 choices.

